# Neuromodulation of striatal D1 cells shapes BOLD fluctuations in anatomically connected thalamic and cortical regions

**DOI:** 10.1101/2022.03.11.483972

**Authors:** Marija Markicevic, Oliver Sturman, Johannes Bohacek, Markus Rudin, Valerio Zerbi, Ben D. Fulcher, Nicole Wenderoth

**Author notes:** V.Z., B.F. and N.W. share last authorship.

## Abstract

Understanding how the brain’s macroscale dynamics are shaped by underlying microscale mechanisms is a key problem in neuroscience. In animal models, we can now investigate this relationship in unprecedented detail by directly manipulating cellular-level properties while measuring the whole-brain response using resting-state fMRI. Here we focused on understanding how blood-oxygen-level-dependent (BOLD) dynamics, measured within a structurally well-defined striato-thalamo-cortical circuit, are shaped by chemogenetically exciting or inhibiting D1 medium spiny neurons (MSNs) of the right dorsomedial striatum (CPdm). We characterize changes in both the BOLD dynamics of individual cortical and subcortical brain areas, and patterns of inter-regional coupling (functional connectivity) between pairs of areas. Using a classification approach based on a large and diverse set of time-series properties, we found that CPdm neuromodulation alters BOLD dynamics within thalamic subregions that project back to dorsomedial striatum. In the cortex, the strongest changes in local dynamics were observed in unimodal regions, i.e., regions that process information from a single sensory modality, while changes in the local dynamics weakened along a putative cortical hierarchical gradient towards transmodal regions. In contrast, a decrease in functional connectivity was observed only for cortico-striatal connections after D1 excitation. Our results provide a comprehensive understanding of how targeted cellular-level manipulations affect local BOLD dynamics at the macroscale, including the role of a circuit’s structural characteristics and hierarchical cortical level in shaping those dynamics. These findings contribute to ongoing attempts to understand the influence of structure–function relationships in shaping inter-regional communication at subcortical and cortical levels.

## Introduction

The brain is a complex network of anatomically connected and perpetually interacting neural components. Perturbing an individual component at the cellular level can alter both the local neural dynamics and patterns of inter-regional communication with the rest of the brain. A large body of work has used functional magnetic resonance imaging at rest (rsfMRI) to investigate interregional functional connectivity (FC) at the macroscale with cellular manipulations (Markicevic et al., 2020; Nakamura et al., 2020; Rocchi et al., 2022; Zerbi et al., 2019) and without (Chuang & Nasrallah, 2017; Gozzi & Schwarz, 2016; Grandjean et al., 2020). But relatively few studies have focused on directly capturing the local dynamical properties of BOLD signal fluctuations *within* brain areas. How microscale alterations shape distributed macroscale dynamical patterns is key to understanding the complex interactions that underly brain dynamics.

One approach to tackling the complex interactions that underly BOLD fluctuations is to understand the influence of structural connections and structure–function relationships in shaping BOLD dynamics. Structural connections constrain functional connectivity, with the strength of this effect varying along a unimodal-to-transmodal cortical gradient in humans: structure–function coupling is higher for unimodal regions that primarily process information from a single sensory modality than for transmodal regions that integrate information across modalities of large scale, polysynaptic circuits (Bazinet et al., 2021; Margulies et al., 2016; Vazquez-Rodriguez et al., 2019). This hierarchical unimodal–transmodal cortical gradient is reflected in microstructural properties, such as cytoarchitecture, size, density and distribution of dendritic cell types, synaptic structure, gene expression, and macroscale functional connectivity in humans and in primates (Burt et al., 2018; Huntenburg et al., 2018). Using an MRI-derived biomarker, the T1-weighted/T2-weighted ratio (T1w/T2w), which is a strong proxy for the structural hierarchy of macaque cortex (Burt et al., 2018), Fulcher et al. (Fulcher et al., 2019) showed that similar gradients also exist in mouse cortex. Indeed, the spatial variation of T1w:T2w across mouse cortex mirrors that of cytoarchitecture, interneuron cell density, long-range axonal connectivity, and gene expression, together ordering cortical areas along a putative functional hierarchy, from unimodal to transmodal (Fulcher et al., 2019). However, it remains unknown whether regional BOLD dynamics also vary along this gradient. Recent work has indicated that local BOLD dynamics are shaped by structural connectivity, with more strongly connected regions exhibiting slower timescales of BOLD dynamics in mouse (Sethi et al., 2017) and human (Fallon et al., 2020). Time-series properties of the BOLD signal also vary with molecular, cellular, and circuit properties of the brain (Gao et al., 2020; Shafiei et al., 2020). While these correlational studies provide an understanding of the relationships between the anatomical properties of brain areas and their BOLD dynamics, direct manipulations are required to provide causal evidence for how cellular-level properties shape BOLD dynamics.

In our previous work, we used rsfMRI with chemogenetic neuromodulation of the somatosensory cortex in mice and measured systematic alterations in BOLD dynamics of the manipulated brain area (Markicevic et al., 2020). Resulting changes to the BOLD time series of local areas were quantified using a machine-learning approach that leverages thousands of time-series features (Fulcher & Jones, 2017; Markicevic et al., 2020). This approach of combining thousands of time series features (including linear autocorrelation, entropy and predictability, and outlier properties), is termed ‘highly comparative time series analysis’ (Fulcher et al., 2013), and provides a data-driven way to assess and interpret changes across a comprehensive candidate set of time-series properties (Fulcher & Jones, 2017). Here we apply this method to regions forming a well-defined striato-thalamo-cortical circuit to understand: i) how a cell-specific perturbation of a striatal region affects the local BOLD dynamics within the manipulated area and within its projection areas, i.e., structurally connected regions in thalamus and cortex; and ii) which structural or functional properties of the projection areas constrain the influence exerted by modulating striatal activity.

To explore this, we chemogenetically either excited or inhibited the activity of D1 medium spiny neurons (MSN) in the right dorsomedial caudate putamen (CPdm, i.e., ‘the input area’ of the striatum (Hintiryan et al., 2016; Lee et al., 2016; Runegaard et al., 2019; Wall et al., 2013)), while acquiring rsfMRI data across the whole brain (Fig. 1A). We investigated how these chemogenetic manipulations shape regional BOLD signal fluctuations (Fig. 1B) in the CPdm-thalamo-cortical loop, evaluating changes across a wide range of time-series properties (Fulcher & Jones, 2017) using a multivariate classification algorithm (Fig. 1A). We found that alterations of D1 MSN activity in the CPdm shape the BOLD dynamics of multiple thalamic and cortical regions. These dynamical changes were strongest for: (i) thalamic subregions which project back to CPdm; and (ii) unimodal rather than transmodal cortical areas, as summarized in Fig. 1B. Our results provide a comprehensive understanding of how targeted cellular-level manipulations affect both local dynamics and interactions at the macroscale and highlight the role of structural circuit characteristics and putative cortical hierarchical level in shaping the dynamics of cortical and subcortical regions.

**Figure 1:**
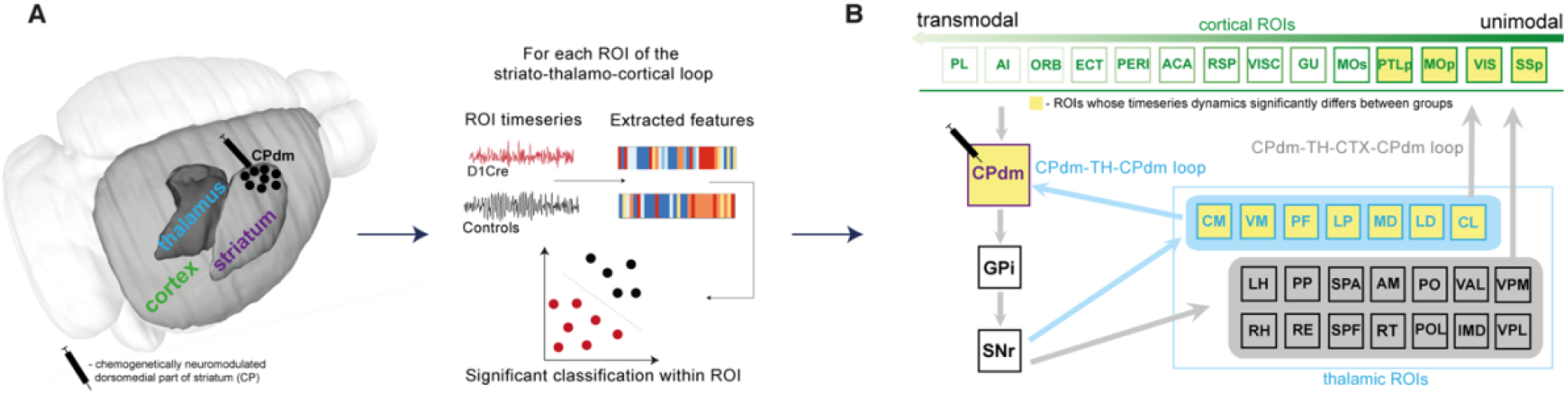
The effect of D1 MSN neuromodulation within dorsomedial striatum was studied within an anatomically defined striato-thalamo-cortical circuit using BOLD time-series properties. **A)** Medium spiny neurons (MSNs) of the right dorsomedial striatum (CPdm) were neuromodulated via DREADDs. Time series were extracted from regions of interests (ROIs) in striatum (CP), thalamus (TH) and cortex (CTX) after the DREADDs were activated by clozapine. BOLD time-series properties were computed using the hctsa feature set (Fulcher & Jones, 2017). For each ROI, a linear support vector machine was used to determine whether neuromodulated animals could be distinguished from controls using time-series features extracted from the BOLD signal. **B)** ROIs within the striato-thalamo-cortical circuit connected to CPdm were identified using mesoscale anatomical atlases. Thalamic areas were subdivided into ROIs that project back to CPdm (light blue box and light blue arrows) versus thalamic ROIs that do not (light grey box). Cortical ROIs (green) were ranked along a previously described multimodal hierarchy (Fulcher et al., 2019). ROIs marked in yellow indicate that neuromodulated versus control animals could be significantly distinguished based on BOLD time-series properties. In cortex, such ROIs are primarily unimodal, while in thalamus such ROIs project back to the neuromodulated CPdm. Full names of abbreviated ROIs: AI - agranular insular area; PL – prelimbic area; ECT|PERI -ectorhinal and perirhinal area; VISC|GU – visceral and gustatory areas; ORB – orbital area; RSP - retrosplenial area; MOp/s - primary and secondary motor cortex; SSp - primary somatosensory cortex; ACA - anterior cingulate area; PTLp - posterior parietal association area; VIS – visual area; CPdm-caudate putamen dorsomedial; SNr – substantia nigra reticular part; GPi – globus pallidus internal; VPL - ventral posterolateral nucleus of thalamus; RE|LH|RH - nucleus of reuniens|lateral habenula|rhomboid nucleus; PO|POL – posterior complex and posterior limiting nucleus of thalamus; SPF|SPA|PP - subparafascicular nucleus subparafascicular area with peripeduncular nucleus of thalamus; IMD -intermediodorsal nucleus of thalamus ; RT - reticular nucleus of thalamus; AM – anteromedial nucleus; CL - central lateral nucleus of thalamus; PF - parafascicular nucleus; VAL|VPM|VPMpc - ventral anterior-lateral complex of the thalamus with ventral posteromedial nucleus of the thalamus and its parvicellular part; MD – mediodorsal nucleus of thalamus; LD - lateral dorsal nucleus of thalamus; LP – lateral posterior nucleus of thalamus; VM|CM – ventral and central medial nuclei of thalamus.

## Results

### Chemogenetic manipulation of D1 MSNs of CPdm alters the motor behavior of animals

Four weeks after surgery, mice underwent a behavioral open-field test after activating the DREADDs with a low dose of clozapine (Fig. 2B). This open-field test was performed as a manipulation check, to test whether activating the DREADDs in CPdm caused behavioral changes as predicted by previous work (Bay Konig et al., 2019; Lee et al., 2016). Exciting D1 MSNs with clozapine increased the number of contraversive rotations and decreased the number of ipsiversive rotations compared to controls who also received clozapine (MANOVA, F_contra_(1,13)=19.7, *p*_contra_=0.001; F_ipsi_(1,13)=8.8, *p*_ipsi_= 0.011; Fig. 2E-F). Exciting D1 MSNs also increased the total distance moved (MANOVA, F(1,13)= 18.2, *p* = 0.001; Fig. 2D). By contrast, inhibiting D1 MSNs of CPdm decreased the number of contraversive rotations, and increased the number of ipsiversive rotations when compared to control mice or to excitatory D1 MSN mice (MANOVA, F_contra_(1,19)=14.7, *p*_contra_=0.001; F_ipsi_(1,19)=66.1, *p*_ipsi_= 1 × 10^−5^; Fig. 2E-F). These results replicate the behavior which is typically observed for unilateral excitation versus inhibition of D1 MSNs (Bay Konig et al., 2019; Lee et al., 2016; Runegaard et al., 2019; Tecuapetla et al., 2014). We also assessed the viral transfection in the CPdm by immunostaining. Figure 2C shows the superimposed viral expression maps of all mice which clearly cover CPdm. These results indicate that our approach successfully modulated D1 MSNs in CPdm.

**Figure 2.**
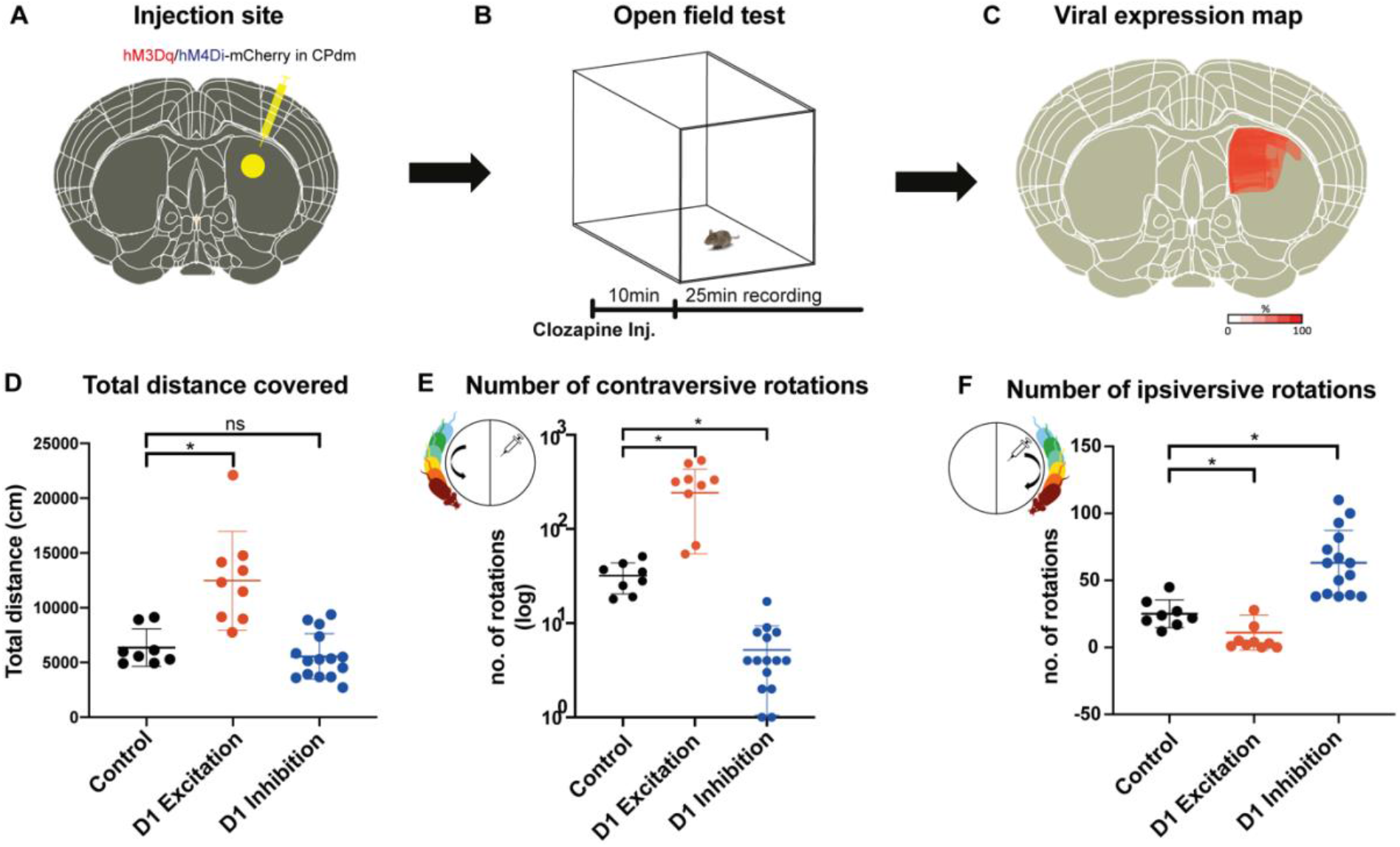
Chemogenetic neuromodulation of D1 MSNs altered the animals’ motor behaviour. **A)** Either DIO-hM3Di-mCherry, DIO-hM4Dq-mCherry or DIO-mCherry (control) virus was injected in the right dorsomedial striatum (CPdm) of D1Cre mice (marked in yellow). **B)** Four weeks after viral injection, an open field test was performed. A 25min recording of mouse behavior was commenced 10min after intraperitoneal clozapine injection. **C)** Qualitative viral expression maps were overlaid for all mice included in the experiment (100% on the scale bar indicates the presence of viral expression in all the mice). **D)** Mice whose D1 MSNs of CPdm were excited and not inhibited, covered significantly more distance when compared to controls (MANOVA, p=0.001). **E)** Exciting D1 MSNs of the right CPdm increased the frequency of contraversive rotations (turning in the direction opposite the injection site as illustrated within the small circle, p=0.001), when compared to controls. On the contrary, inhibiting D1 MSNs of the right CPdm decreased contraversive rotations relative to controls (p=0.001). F) Compared to controls, the number of ipsiversive rotations (turning in the same direction as the injection site, as illustrated within the small circle) significantly decreased when D1 MSNs of the CPdm were excited (p=0.011), while rotations significantly increased when D1 MSNs were inhibited (p=1.3*10^−5^).

### Altered dynamics of neuromodulated CP subregion and its anatomically adjacent regions

Seven days after behavioral testing, mice were lightly anesthetized and spontaneous brain activity was measured via rsfMRI before and after activating the DREADDs with clozapine. Since we neuromodulated D1 MSNs in the intermediate portion of the right dorsomedial part of CP (CPdm), we were specifically interested in whether this target area or other right striatal subregions would be affected. As described in Methods, we trained separate time-series classifiers for each of the 29 CP subregions to determine whether changes in the properties of BOLD time series within individual brain areas were caused by D1 excitation versus no modulation in control mice. We repeated this exploratory analysis to assess D1 inhibition versus no modulation in control mice.

Balanced classification accuracies (and permutation-test *p*-values) for each of the 29 CP subregions are summarized in Figs 3A-B (see Table S1 for full results). Regions varied widely in their classification accuracy, ranging from chance-level to >85%, for both group comparisons (D1 control vs D1 excitation and D1 control vs D1 inhibition). High classification accuracy (>75%) was obtained when comparing BOLD dynamics within the injection site (CPi,dm,cd) between excitation/inhibition of D1 MSNs and control mice (all *p*_uncorr_ < 0.03, Suppl. Table S1), although this was not statistically significant after the correction for multiple comparisons. In addition to the injection site, we also found changes in dynamics of anatomically adjacent CP sub-areas in the intermediate-ventral portion or the rostral-dorsomedial portion of the CP (all *p*_uncorr_ < 0.05, Table S1), which also survived correction for multiple comparisons for D1 excitation group (Fig. 3A). In summary, our exploratory analyses indicate high classification accuracy in DREADD targeted areas and anatomically adjacent CP sub-areas.

**Figure 3.**
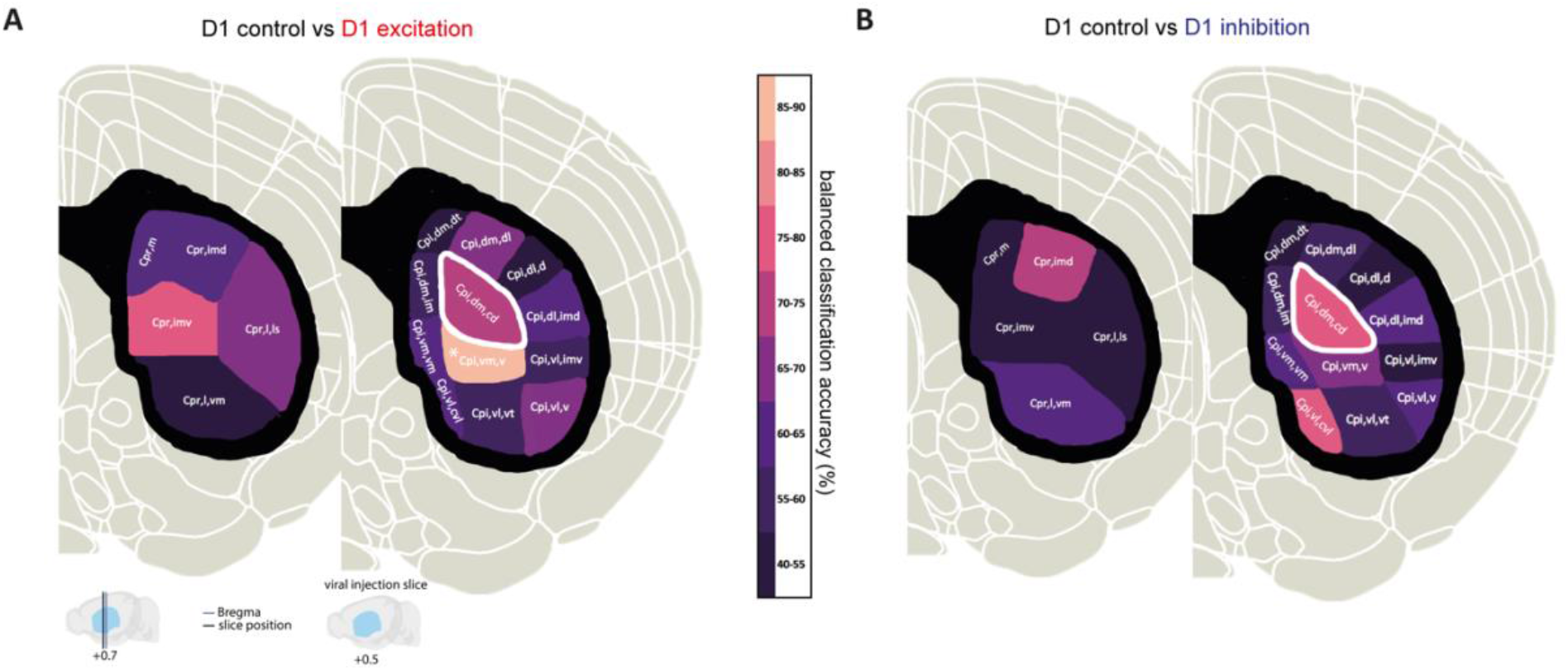
Alterations in local BOLD dynamics within the striatum. **A) - B)** Coronal slices of the mouse brain with color-coded balanced classification accuracy results for CP subregions obtained from the Mouse Cortico-Striatal Projectome (Hintiryan et al., 2016) for D1 control vs D1 excitation groups (A) and D1 control vs D1 inhibition (B), respectively. Region circled with the white border is the exact DREADD injection site. * indicates FDR corrected significance (p < 0.05).

### Changes in BOLD dynamics in thalamic sub-regions that form anatomically closed loops with CPdm

We next sought to understand whether activating the DREADDs in CPdm affects the local BOLD dynamics of anatomically connected thalamic nuclei. In each thalamic ROI, we applied a classification analysis to the MSN D1 excitation versus control group and the MSN D1 inhibition versus control group. We found statistically significant classification accuracies (*p*_corr_ < 0.05, permutation test) for multiple thalamic ROIs: parafascicular nuclei (PF), lateral dorsal (LD), lateral posterior (LP), mediodorsal (MD), and ventral and central medial (VM|CM) nuclei of thalamus. Interestingly, the largest effects were observed for those thalamic ROIs that have a specific reciprocal anatomical connection with CPdm (Fig. 4A; Fig. S1A-B), i.e., they receive projections from CPdm via GP/SN and project back to CPdm. Indeed, ROIs with reciprocal anatomical connections to CPdm displayed higher classification accuracies than other thalamic ROIs for both D1 excitation vs D1 control (two-tailed Mann-Whitney U test, U(8,6)=0, z=-3.10, *p*=7×10^−4^, Fig. 4B), and D1 inhibition vs D1 control (two-tailed Mann-Whitney U test, U(8,6)=8, z=-2.07, *p*=0.04, Fig. 4C). In summary, we found that modulating D1 MSN activity alters BOLD signal dynamics in downstream thalamic regions that form strong closed-loop anatomical connections with the neuromodulated CPdm.

**Figure 4:**
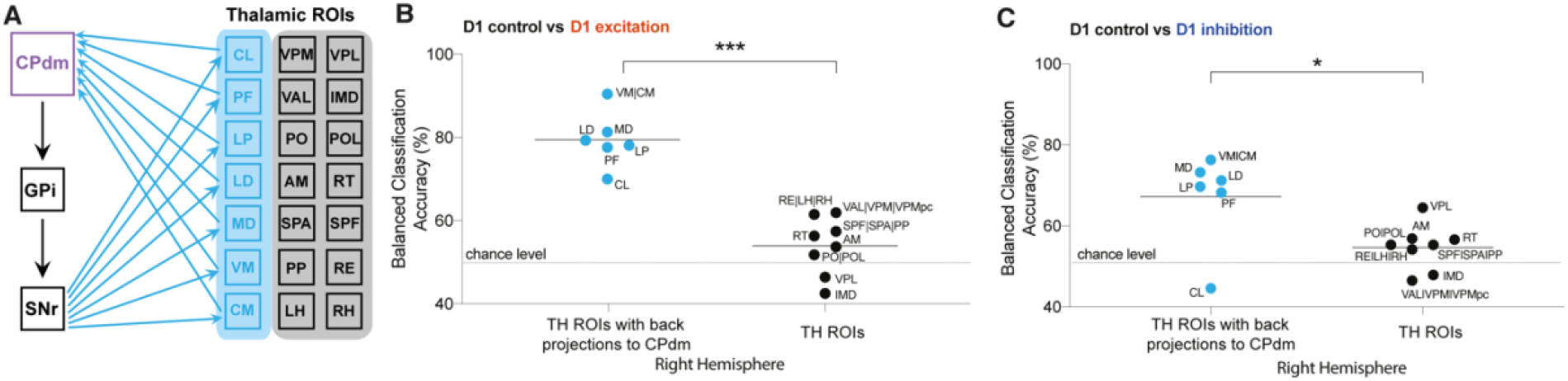
Altered BOLD dynamics in thalamic regions forming anatomical loops with dorsomedial striatum A) Figure illustrating thalamic (TH) subregions (light blue) that project back to dorsomedial striatum. **B)** Comparison of balanced classification accuracies for D1 control vs D1 excitation group between thalamic regions projecting back to the modulated site CPdm (blue dots) and the rest of the thalamic ROIs which constitute striato-thalamo-cortical circuit (two-tailed Mann-Whitney U test; p = 7×10^−4^). Thalamic regions which project back to CPdm have significantly higher balanced classification accuracies as compared to the other thalamic ROIs. **C)** Similar to B but for D1 control vs D1 inhibitory group comparison (two-tailed Mann-Whitney U test; p = 0.04). Full names of abbreviated ROIs in Fig. 1 caption.

### Cortical regions with altered BOLD dynamics are primarily unimodal

Next we evaluated the classification accuracy among cortical ROIs during D1 MSN CPdm neuromodulation. Using our feature-based time-series classification approach, we assessed changes in the BOLD dynamics of individual cortical areas for D1 excitation versus no modulation in control mice. Significant changes in BOLD dynamics (*p*_corr_ < 0.05, permutation test) were observed for four cortical ROIs: SSp – primary somatosensory area, VIS – visual area, MOp – primary motor area and PTLp – posterior parietal association area (Fig. 5A). Repeating the analysis for D1 inhibition versus control resulted in two significant regions: PTLp and SSp (Fig. 5B).

**Figure 5:**
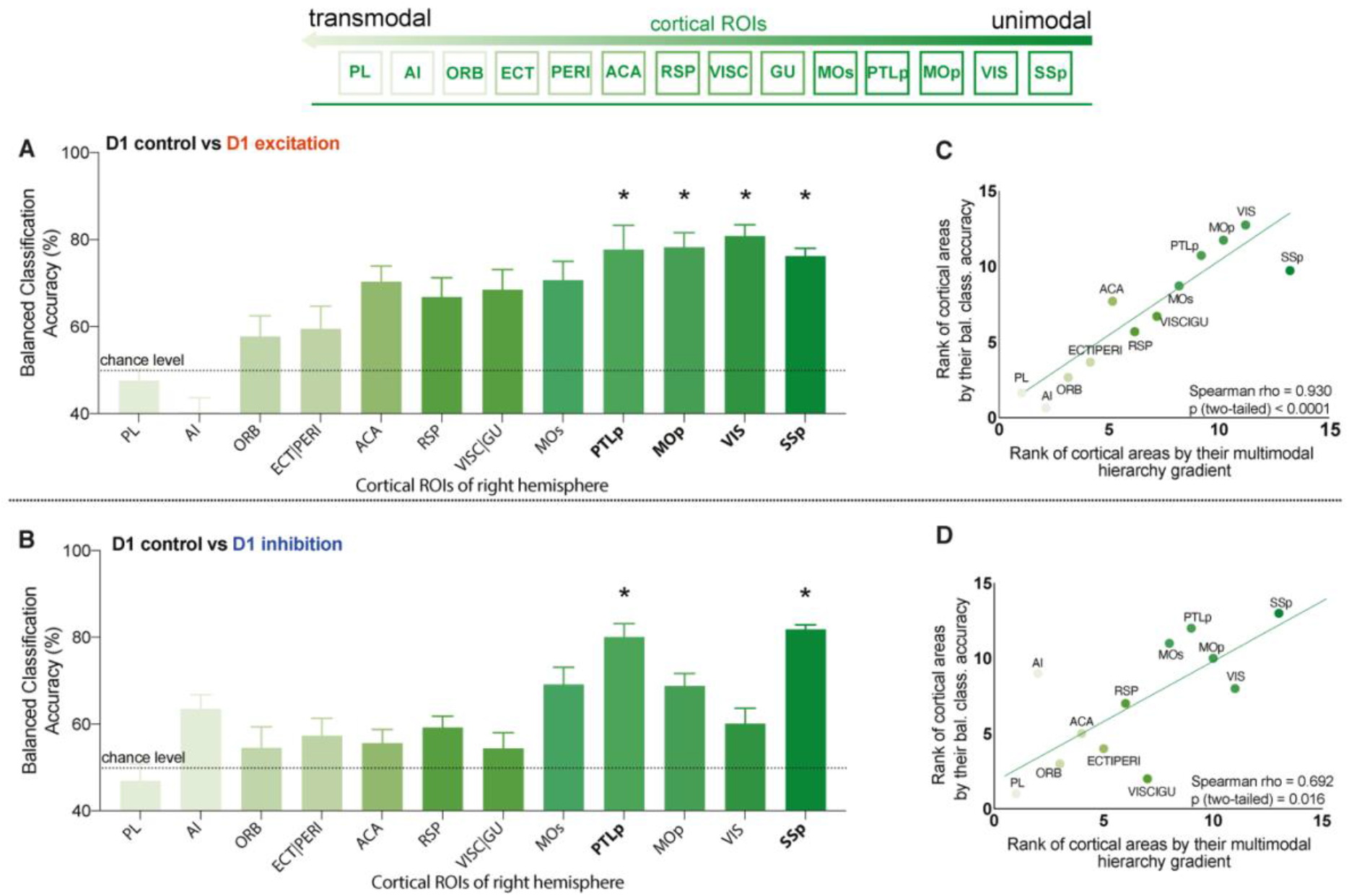
Cortical ROIs with altered BOLD dynamics after D1 MSN modulation are primarily unimodal. **A) – B)** Balanced classification accuracy (%) results for each cortical brain region shown for: **A)** D1 control versus D1 excitation; **B)** D1 control versus D1 inhibition. Regions are listed following a previously described hierarchical ordering of cortical areas (Fulcher et al., 2019) that putatively orders regions along a unimodal-to-transmodal axis (illustrated above the two figures). An FDR corrected significant difference in BOLD dynamics between the two groups is depicted with bold ROI abbreviations and an asterisk (*) (pcorr<0.05). **C)** Spearman correlation (rho = 0.93, p=1×10^−4^) between cortical areas ranked by their balanced classification accuracy and their multimodal hierarchy gradient for D1 control vs D1 excitation group comparison. **D)** Same as C but for D1 control vs D1 inhibition group comparison (rho=0.69, p=0.02). Full names of abbreviated ROIs in Fig. 1.

We next aimed to understand whether the observed differences in response to D1 MSN manipulation across cortical areas could be related to other sources of heterogeneity across cortical areas. In particular, recent work has shown a key spatial dimension of heterogeneity across mouse cortical areas, in which gene-expression patterns, cell-type densities, and other structural properties follow a common, and putatively hierarchical variation from primary unimodal areas to integrative transmodal association areas (Fulcher et al., 2019; Huntenburg et al., 2021). Combining information from normalized measures of cell densities, cytoarchitecture, T1w:T2w, and brain-relevant gene-expression markers, Fulcher et al. (2019) used principal components analysis to extract an ordering of cortical areas from transmodal (lowest rank) to unimodal (highest rank). We assessed whether this hierarchical ordering of cortical areas was related to the variation in their dynamical response to D1 MSN neuromodulation in CPdm. The relationship between cortical ROIs ranked according to the Fulcher et al. multimodal hierarchy, and by our balanced accuracy, is shown in Figs 5A-B. We find a strong positive correlation between the two measurements for both types of modulation: i) D1 excitation vs D1 control (Spearman’s ρ = 0.93, *p* = 0.0001, Fig. 5C); and ii) D1 inhibition vs D1 control (Spearman’s ρ = 0.69, *p* = 0.02, Fig. 5D). Our results indicate that D1 MSN modulation alters BOLD dynamics more strongly in unimodal cortical regions than in transmodal cortical regions.

### Common time-series properties drive successful classification among thalamic regions

Our results demonstrate that changes in macroscopic BOLD dynamics in response to D1 MSN neuromodulation of CPdm can be assessed using a feature-based time-series classification approach. But which properties of BOLD time series are specific to each manipulation? To answer this, we investigated each cortical (SSp, PTLp, MOp and VIS) and thalamic (PF, LP, LD, MD and VM|CM) ROI with significantly altered dynamics after D1 excitation. We computed the signed Mann-Whitney rank-sum statistic and associated *p*-value, corrected across all features in each cortical and thalamic ROIs (see Methods). We found individual significant features in the LD (284 significant features) and VM|CM (344 significant features). Individual significant features were further inspected, in order to find a common signature across regions. In both regions, significant features belonged to autocorrelation and frequency-based statistics, displaying increased simple linear autocorrelations at short time lags (e.g., autocorrelation at lag 1, AC(1), Fig. 6A-B), and an increase in low-frequency spectral power. This is consistent with the widespread use of linear correlation-based statistics, like fALFF, to analyze fMRI time series (Fallon et al., 2020; Sethi et al., 2017; Shafiei et al., 2020; Shinn et al., 2021; Zou et al., 2008).

**Figure 6:**
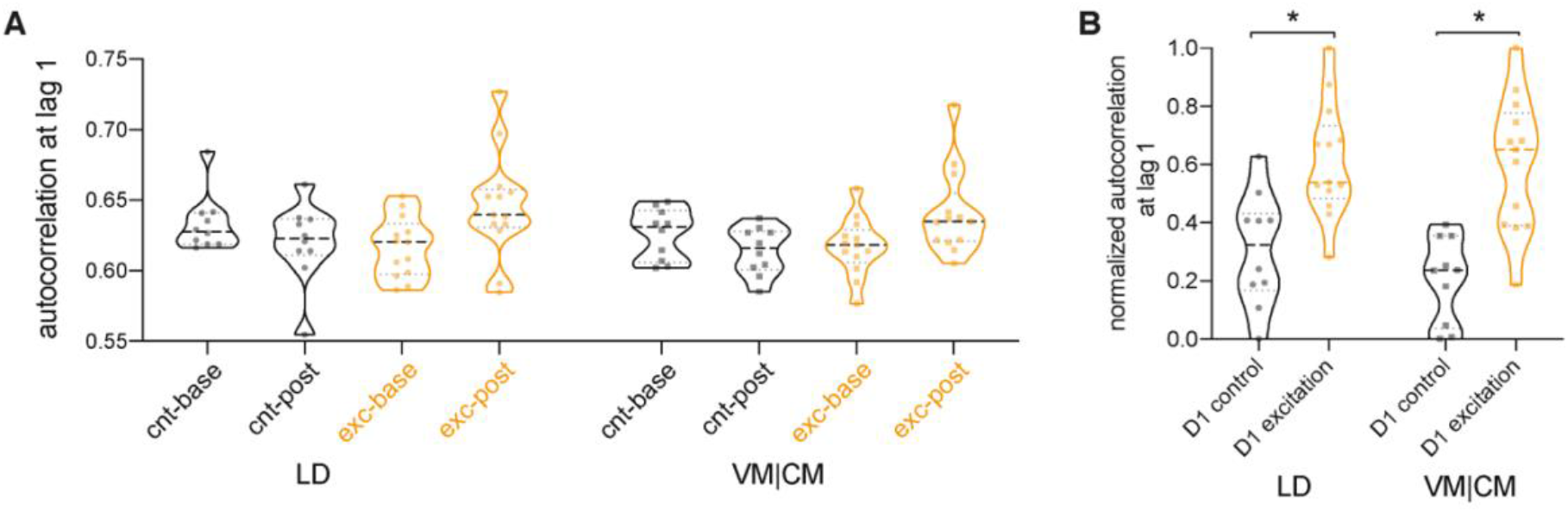
D1 MSN excitation led to slower and more autocorrelated fluctuations of the timeseries signal in thalamic regions. **A)** Autocorrelation at lag 1 shown for baseline and post clozapine controls (cnt-base, cnt-post, respectively) and excitatory groups (exc-base, exc-post) for lateral-dorsal (LD) and ventral-, central-medial (VM|CM) thalamic regions. Increase in autocorrelation is observed during a post clozapine period in the excitatory group, for both brain regions (LD and VM|CM). **B)** Increase in autocorrelation is observed in normalized within-group baseline-corrected data. * indicates a between groups FDR corrected significant difference i.e., p_LD_=0.034; p_VM|CM_=0.009.

For other ROIs, no individual features differed significantly between D1 excitation and control conditions (after multiple hypothesis correction), indicating that the classification results were driven by combined contributions across multiple different features. However, an exploratory analysis revealed that individual time-series features with the strongest discriminative ability in other ROIs (SSp, PTLp, Mop, VIS, PF, LP and MD) were qualitatively similar to those identified for LD and VM|CM, i.e., broadly measuring increased autocorrelation of the BOLD signal following D1 excitation. Our exploratory analysis did not find any individually significant features following D1 inhibition.

### Multiple cortical regions display changes in functional connectivity after CPdm neuromodulation

Since excitation and inhibition of D1 MSNs of CPdm significantly alters the local dynamics of remote brain regions in the striato-thalamo-cortical circuit, we next examined whether their pairwise coupling, measured as functional connectivity (FC) using linear Pearson correlations, was also affected (Fig. 7). We assessed how injecting clozapine changes FC between CPdm and each of the anatomically connected ROIs.

**Figure 7:**
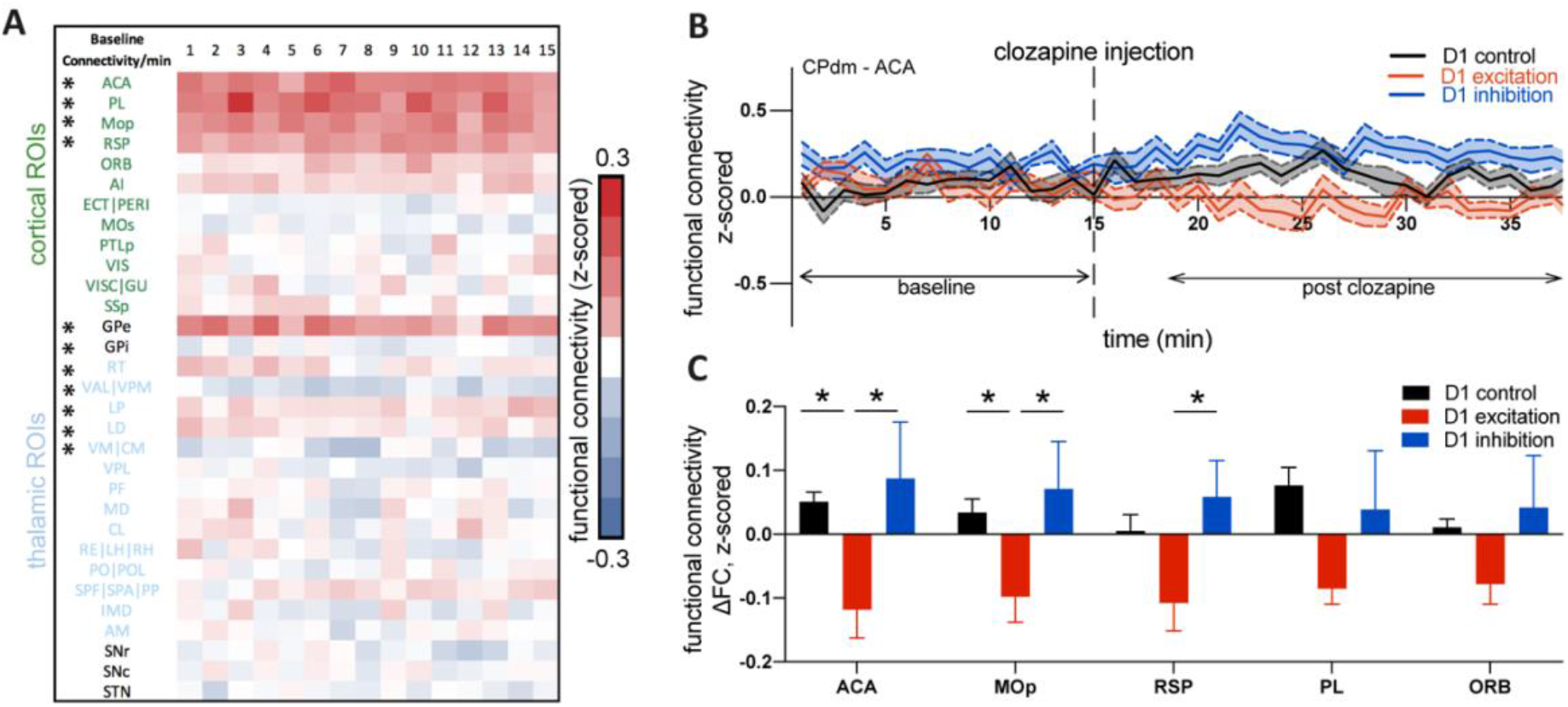
Cortico-striatal changes in functional connectivity upon D1 MSN neuromodulation. **A)** Baseline functional connectivity (FC) calculated per minute between CPdm and the rest of ROIs that form the striato-thalamo-cortical loop, combining animals from all three groups involved in the experiment. One-sample Wilcoxon test was performed and * indicates that statistical significance was reached after FDR correction for multiple comparisons (p<0.01). **B)** Functional connectivity calculated by minute for D1 controls, D1 excitation and D1 inhibition groups between CPdm and ACA (an example ROI). Dashed line at 15min indicates clozapine injection i.e., activation of the DREADD, which is followed by the changes in FC for D1 excitation and D1 inhibition groups. The baseline (first 15min before clozapine injection) and post clozapine periods (from 19 – 34 min) used for the delta analysis (Fig. 7C) are indicates. **C)** Functional connectivity at ΔFC (post clozapine – baseline) for cortical regions that displayed differences during pairwise comparisons (permutation test, p_uncorr_ < 0.05). * indicates that statistical significance was reached after FDR correction on all pairwise comparisons of all 3 groups.

First, animals from all three groups were pooled and baseline FC (i.e., before the DREADDs were activated with clozapine) was calculated. One-sample Wilcoxon tests revealed that ACA, PL, MOp, RSP, GPe, RT, LP and LD were positively correlated with CPdm, while GPi, VAL|VPM and VM|CM were negatively correlated (*p*_corr_ < 0.01, FDR corrected, Fig. 7A). Next, we investigated how FC changed from baseline to the post-clozapine period, for each of the three groups and each ROI. For example, FC between CPdm and ACA (Fig. 7B) was reduced when D1 MSNs were excited (Fig. 7B, red trace), while it was increased when D1 MSNs were inhibited (Fig. 7B, blue trace), and it remained largely unchanged in the control group (Fig. 7B, black trace).

Direct comparison between D1 excitation and controls indicated a significant decrease in FC connectivity after D1 excitation between CPdm – ACA and CPdm – MOp connections (*p*_corr_ < 0.05, FDR-corrected, Fig. 7C). Additionally, when we compared whether exciting versus inhibiting D1 MSNs differentially affects CPdm connectivity, we saw that excitation typically reduced FC, while there was a trend towards increased FC during inhibition (Fig. 7C). This was statistically significant for FC changes between CPdm and ACA, and RSP and MOp (*p*_corr_ < 0.05, FDR-corrected). We also performed an analysis of DREADD-induced FC changes among all ROIs of striato-thalamo-cortical circuit, including thalamo-striatal and thalamo-cortical connections, but found no significant connections (*p*_corr_ < 0.05, FDR-corrected; data not shown). Overall, our FC results indicate a significant decrease in FC between CPdm and three cortical nodes, when D1 MSNs were excited and compared to either controls or D1 inhibition.

## Discussion

Understanding how the brain’s macroscale dynamics are shaped by underlying microscale mechanisms is a key problem in neuroscience, that cannot be addressed directly using conventional correlational studies. Here, we used an anatomically well-defined striato-thalamo-cortical circuit to investigate how cell-specific chemogenetic excitation or inhibition of D1 MSNs of right dorsomedial striatum affects regional BOLD dynamics and pairwise coupling. Using a classification approach based on a large and diverse set of time-series properties from the *hctsa* toolbox (Fulcher & Jones, 2017), we found that CPdm neuromodulation alters BOLD signal dynamics in anatomically connected sub-areas of thalamus and cortex. Importantly, the regional changes in BOLD dynamics were stronger for: i) thalamic regions with reciprocal, structural connectivity with CPdm; and ii) cortical areas low in a putative cortical hierarchy. We found that D1 excitation increases the autocorrelation and low-frequency power of the BOLD signal, showing how neuromodulation shapes the temporal autocorrelation of distributed but connected regions at the macroscale. We also assessed functional connectivity between the neuromodulated dorsomedial striatum and areas of the connected striato-thalamo-cortical circuit, but only observed decreased striato-cortical functional connectivity after D1 MSN excitation. Taken together, we show that fMRI time-series properties can be used to study how the dynamics within macroscopic brain regions are affected by targeted cellular-level neural manipulations. Our results suggest that regional BOLD dynamics are at least partially shaped by the structural connections of a circuit, as well as variation along gradients in cortical architecture, with unimodal regions responding more strongly to perturbations from dorsomedial CP. Our work provides a comprehensive understanding of how targeted cellular-level manipulations affect both local dynamics and interactions at the macroscale and demonstrate the influence of structural characteristics of a circuit and putative cortical hierarchy in shaping those dynamics.

### Altered BOLD dynamics of a targeted CPdm and its anatomically adjacent CP subareas

Our results confirm that after activating the DREADDs in CPdm, BOLD time-series properties change in the targeted striatal sub-areas. We also observed substantial modulation of BOLD dynamics in anatomically adjacent CP subregions, located either ventrally or rostrally to the injection area, CPdm (Fig. 3). While we cannot exclude the possibility that the injected virus spread to neighboring CP subareas, it is unlikely that virus-spread alone explains the robust neuromodulatory effects which we observed at the group level. An alternative explanation is that our results reflect intra-striatal communication between the identified CP sub-areas. The striatum is the main input structure of the basal ganglia and receives glutamatergic inputs from cortical, thalamic and limbic regions combined with dopaminergic input from the midbrain, thus acting as an integrative hub that assists in the selection of appropriate behaviors through its output to downstream basal ganglia nuclei (Reig & Silberberg, 2014). This integrative function of the striatum is thought to happen at the level of the local intra-striatal circuitry (Burke et al., 2017). Intra-striatal connectivity involves synaptic connections between MSNs and striatal interneurons. However, there is also evidence of functional neuronal ensembles, which are formed by widely distributed MSNs throughout the striatum, not necessarily in close proximity to each other (Burke et al., 2017). Specifically, *in vivo* electrophysiological recordings across a large fraction of striatum along the CPdm and CP-ventral axis, revealed that ensembles of MSN, which fired in a highly correlated manner, were formed throughout the recorded striatal subdivisions (Bakhurin et al., 2016). Activity in these MSNs ensembles was not only highly correlated when the animal executed a task but also during the resting-state period (Bakhurin et al., 2016). Moreover, cortico-striatal projections that project to distinct intermediate parts of CP were found to extend their projections to the rostral part of CP, indicating another possible avenue for intra-striatal communication mediated by cortical inputs (Hintiryan et al., 2016). Based on these findings it is tempting to speculate that chemogenetic neuromodulation of D1 MSNs altered not only the BOLD dynamics of the target CPdm, but also of intra-striatally connected subareas. Future experiments tackling structural and functional intra-striatal communication and their relationship could further our understanding of the changes observed at the level of BOLD dynamics.

### Altered BOLD dynamics in thalamic regions mediated by direct anatomical projections with CPdm

The thalamus is a highly heterogeneous structure, forming bidirectional connections with visual, limbic, associative, sensory and motor regions of the cortex as well as striatum. Extensive research has focused on anatomically mapping the striatothalamic and thalamostriatal projections (Alloway et al., 2017; Antal et al., 2014; Bubb et al., 2017; Diaz-Hernandez et al., 2018; El-Boustani et al., 2020; Elena Erro et al., 2002; Guo et al., 2015; Hunnicutt et al., 2016; Kamishina et al., 2008; Lee et al., 2020; Linke et al., 2000; Mandelbaum et al., 2019; Namboodiri et al., 2016; Parker et al., 2016; Perry & Mitchell, 2019; Smith et al., 2004; Van der Werf et al., 2002; Vertes et al., 2015; Wall et al., 2013; Wang et al., 2006), which enabled us to further distinguish thalamic ROIs based on their projections to/from the neuromodulated CPdm. Here we show that after D1 MSN excitation and inhibition, those thalamic ROIs that form anatomically closed loops with CPdm display the strongest alteration in BOLD dynamics. A correlational analysis of the relationship between structural connectivity and BOLD dynamics from Sethi and colleagues (Sethi et al., 2017) suggested that regional BOLD dynamics are shaped by the strength of incoming axonal projections. Our CPdm manipulations that resulted in BOLD signal perturbations of only thalamic regions with anatomical projections originating in the CPdm provide causal evidence in support of this hypothesis. Our findings contribute to ongoing attempts to understand the influence of structural connections, as a function of their strength, in shaping inter-regional communication among cortical and subcortical regions. Complementing traditional patterns of functional connectivity in the resting state, the empirical results presented here provide important new understanding of how the brain responds to perturbations, information that will help develop, refine, and validate mathematical models of the brain’s distributed dynamics.

### Cortical regions with altered BOLD dynamics are primarily unimodal

Out of twelve cortical regions included in the striato-thalamo-cortical circuit, four exhibited significantly altered BOLD dynamics after D1 MSN excitation and two after D1 MSN inhibition. This is remarkable because it provides further evidence that modulating D1 MSN activity in the CPdm causes changes in BOLD dynamics which propagate through poly-synaptically connected circuits. There was substantial variation in how strongly the BOLD dynamics of cortical regions changed in response to neuromodulating striatal D1 MSN cells. Our results indicate that this variation follows a recently described hierarchical gradient (Fulcher et al., 2019): a spatial variation across the cortex that is broadly shared by diverse measurements of cortical microstructure including cytoarchitectural types, neural cell densities, T1w/T2w, axonal input strength, and the expression of brain-relevant genes. Here we show that the influence of D1 neuromodulation of CPdm on the dynamics of cortical regions strikingly mirrors this hierarchical variation, with putatively unimodal areas lower in the hierarchy being more sensitive to D1 neuromodulation of CPdm than areas higher in the hierarchy.

Compared to transmodal regions, which are responsible for integrating large numbers of diverse inputs, unimodal regions generally have fewer interareal inputs and a less flexible (more ‘hard-wired’) function (Huntenburg et al., 2018). As a result, perturbing the input to a unimodal region (such as that from CPdm) may constitute a greater fraction of their total input relative to a transmodal region (which receives a greater number of inputs), and therefore yield a greater response. This picture is consistent with the hypothesis that the aggregate strength of axonal inputs plays a key role in shaping the BOLD dynamics of an area (Sethi et al., 2017). Alternatively, the underlying biological mechanism may be more specific, and could correspond to any of the many biologically plausible cortical properties that vary hierarchically across the mouse cortex, e.g., the response may be shaped by the hierarchical variation of one or a combination of cell densities and synaptic receptors (Fulcher et al., 2019). Our results thus suggest a range of candidate mechanisms, which future investigations may distinguish, towards a better understanding how interareal communication shapes the dynamics of individual cortical areas.

### D1 MSN excitation leads to slower and more autocorrelated fluctuations of the timeseries signal

An increase in autocorrelation of the BOLD time series was observed for two significant thalamic regions following D1 excitation (measured by properties of the autocorrelation function and Fourier power spectrum). These results are in line with the widespread fMRI literature investigating fMRI timeseries using linear correlation-based statistics (Fallon et al., 2020; Nakamura et al., 2020; Sethi et al., 2017; Shinn et al., 2021; Zou et al., 2008). Specifically, Nakamura and colleagues (Nakamura et al., 2020) detected an increase in the fractional amplitude of low-frequency fluctuations (fALFF) after excitation of D1 MSNs in the dorsal CP, which is in line with our own results that revealed an increase in low-frequency power, albeit in thalamic regions. Moreover, Shafiei and colleagues (Shafiei et al., 2020) report that regional differences in temporal autocorrelation of the human cortex are correlated to molecular and cellular properties of cortical regions. Following this notion, our results provide causal evidence that cellular excitation of D1 MSNs shapes the temporal autocorrelation of anatomically connected regions at the macroscale.

The mechanisms through which D1 excitation increases the autocorrelation of BOLD dynamics within regions of striato-thalamo-cortical circuitry remain unexplored. Assessment of neuronal spike-count autocorrelation has shown that different regions exhibit varied timescales of fluctuations in neuronal population activity i.e., magnitude of autocorrelation (Churchland et al., 2011; Murray et al., 2014), which varied in the presence of a stimulus or a task (Nougaret et al., 2021; Runyan et al., 2017; Wasmuht et al., 2018). Using calcium imaging, increased autocorrelation has been observed in parietal cortex neurons following an auditory decision-making task in mice (Runyan et al., 2017), showing that autocorrelation is a statistical property that enables information coding by neuronal populations over different timescales. The variability of the magnitude of the autocorrelation of intrinsic neural signal was also demonstrated using other neuroimaging modalities such as resting-state fMRI data and human intracranial recordings (Gao et al., 2020; Stephens et al., 2013; Watanabe et al., 2019). Based on this, we speculate that D1 CPdm neuromodulation alters neuronal autocorrelation at the population level, shaping the dynamics of the fMRI signal, which is also characterized by an increase in autocorrelated fluctuations. This could mean autocorrelation is a statistical property that is informative of changes observed across different spatial scales of a dynamical system i.e., from micro-to macro-scales. It is also possible that, due to the characteristic spectral profile of the hemodynamic response to neural activity, alterations in linear correlation properties of the BOLD signal reflect changes in the relative amount of neural activity following D1 CPdm neuromodulation (the signal:noise ratio). For a detailed mechanistic understanding of changes in autocorrelation across different spatial scales, future multimodal experiments combining fMRI with invasive measurements, such as calcium imaging in animal models, as well as computational modelling are necessary.

### Multiple cortical regions display changes in functional connectivity after CPdm neuromodulation

We demonstrate differential modulation of striato-cortical functional connectivity between CPdm and the rest of the striato-thalamo-cortical regions. Nakamura and colleagues (Nakamura et al., 2020) have performed electrophysiological recordings in both dorsal striatum and motor cortex before and after D1 MSN excitation. Results indicate that neuromodulation increased delta power in dorsal striatum but not in motor cortex. Since slow oscillations of delta-band fluctuations contribute to functional connectivity (Wang et al., 2012) and are characterized by brain-wide synchrony (Pan et al., 2013; Uhlhaas et al., 2010), we speculate that differences in low-frequency power changes between the neuromodulated CPdm and cortex contribute to our observations of decreased striato-cortical functional connectivity after exciting D1 MSNs. Moreover, recent findings illustrate that low power coherence between chemogenetically inhibited areas and remote non-neuromodulated areas also contributes to shaping functional connectivity between those regions (Rocchi et al., 2022) . This indicates that more comprehensive analysis combining electrophysiology with fMRI could provide clearer insights into the mechanisms behind observed changes in functional connectivity.

### Limitations and interpretational issues

The data-driven approach of classifying BOLD time series using a comprehensive range of time-series analysis methods allows us to capture diverse types of changes in dynamical properties. While the resulting classification models, involving large numbers of extracted features, can be difficult to constrain on limited sample sizes, given the relative complexity of the feature space we used a relatively simple (linear SVM) classification model. Using this approach, we report strong and highly significant cross-validated classification accuracies here. While the combined information from thousands of features provided strong classification performance, the need to statistically correct across many thousands of features limited our ability to pinpoint the contribution of individual features in most brain areas. This may be overcome with a larger sample size, which may allow us to determine the combinations of time-series features that drive the changes to BOLD dynamics or, alternatively, by comparing a more constrained set of features.

We did not find evidence of altered BOLD dynamics of the basal ganglia structures downstream from CP, namely GPi/e, SNr/c and STN by exciting or inhibiting D1 MSNs. While this is at odds with conventional models of the basal ganglia, some studies indicate that after D1 excitation or D1 inhibition within the dorsal striatum, the firing rates of these downstream structures change in a non-uniform manner i.e., with partial firing rate increases and partial decreases within the single structure (Freeze et al., 2013; Kravitz et al., 2010; Lee et al., 2016; Tecuapetla et al., 2014). In addition, a recent tracer study showed that projections from CP to the output nuclei GPi and SNr are topographically segregated so that CPdm projects only to a specific anatomically defined section of GPi/SNr (Lee et al., 2020). If only a small fraction of an already-small nucleus is affected by CPdm neuromodulation, the spatial resolution of the BOLD signal is insufficient. This would explain why the BOLD dynamics of regions downstream of are only moderately affected by D1 MSN excitation or inhibition.

## Conclusions

In summary, we characterized the influence of local cellular perturbations on macroscale BOLD dynamics for each region within an anatomically connected system. Our results indicate that alterations of BOLD dynamics are shaped by: i) anatomical connectivity of thalamic subregions with CPdm; and ii) the relative position of cortical regions along a putative cortical hierarchy. Our results provide a comprehensive understanding of how targeted cellular-level manipulations affect both local dynamics and interactions at the macroscale, revealing the influence of structural characteristics of a circuit and hierarchical cortical gradients in shaping those dynamics. Furthermore, we provide causal evidence into how regional dynamics change after neuromodulation shaping the temporal autocorrelation of distributed but connected regions at the macroscale. Our findings contribute to ongoing attempts to understand the influence of structure–function relationships in shaping inter-regional communication at subcortical and cortical levels. The results reported here could help in developing, refining, and validating dynamical models for the brain’s distributed dynamics.

## Materials and Methods

All experiments and procedures were conducted following the Swiss federal Ordinance for animal experimentation and approved by Zurich cantonal Veterinary Office (ZH238/15 and ZH062/18). House inbred BAC-mediated transgenic mouse line from GENSAT (BAC-Cre Drd1a-262 – D1Cre) (Gong et al., 2003) was used in this study. All D1Cre mice were kept in standard housing under 12h light/dark cycle with food and water provided *ad libitum* throughout the whole experiment. A total of 38 mice were used in the experiment, aged 16.2 ± 2.8 weeks and weighing 24.9 ± 3.3 g at the day of the surgery. Prior to surgery, the mice were randomly assigned to any of the 3 groups i.e., D1 controls, D1 inhibition or D1 excitation. Following the 3R measures (https://www.swiss3rcc.org/en/3rs-resources/what-are-the-3rs) and based on our previous experience with DREADD experiments (Markicevic et al., 2020; Zerbi et al., 2019), our sample size goal per group was set to an approximate n=15 for D1 inhibition D1 excitation and n=10 for D1 controls.

### Stereotactic transfection procedure

Each mouse was initially anesthetized using a mixture of midazolam (5 mg/mL; Sintetica, Switzerland), fentanyl (50 mg/mL; Actavis AG, Switzerland) and medetomidine (1 mg/mL; Orion Pharma, Finland). After anesthesia induction, mice were placed on a heating pad and the temperature was kept at 35°C (Harvard Apparatus, USA). Following shaving and cleaning, an incision along the midline of scalp was made. The intermediate portion of the right dorsomedial caudate-putamen (CPdm) was targeted at the coordinates of +0.5 mm AP (anterior-posterior), -1.5 mm ML (medio-lateral) and -3.0 mm DV (dorso-ventral) relative to the Bregma using a drill and microinjection robot (Neurostar, Germany) with a 10 μl NanoFil syringe and 34Ga bevelled needle (World Precision Instruments, Germany). 950 nl of double-floxed inverted (DIO) recombinant AAV8 virus was used to express either hM3Dq-mCherry (excitatory DREADD, *n* = 13, 8 females) or hM4Di-mCherry (inhibitory DREADD, *n* = 15, 7 females) or mCherry (control, *n* = 10, 6 females). The virus was injected at the rate of 0.06 ul/min and provided by Viral Vector Core Facility of the Neuroscience Centre Zurich (http://www.vvf.uzh.ch/en.html). After injection, the needle was left in place for 10 min and then slowly withdrawn. Subsequently, mice were given an anesthesia antidote consisting of temgesic (0.3 mg/mL; Reckitt Benckiser AG, Switzerland), annexate (0.1 mg/mL; Swissmedic, Switzerland) and antisedan (0.1 mg/mL; Orion Pharma, Finland) and left to fully recover. Following the surgery, ketoprofen (10 mg/kg; Richter Pharma AG, Austria) was subcutaneously injected daily for at least 3 days to reduce any post-operative pain. Animals were given 3-4 weeks to fully recover from the surgery and to allow for expression of the transgene prior to the scanning session. The viral expression map showing the distribution of the viral expression of all mice included in this study is shown in Fig. 2C.

### Behavioral open field test

A custom-made box (50 × 50 × 50 cm) consisting of light grey walls and a floor was designed and placed in a room with a homogenously spread light source. Each mouse spent 5 min exploring the box before the start of the experiment. To activate the DREADD, clozapine was intraperitoneally injected at a dose of 30 μg/kg, 10 min before the start of the recording. A total of 32 mice underwent an open field behavioural test and each recording lasted 25 min. Data was analyzed using EthoVision XT14 (Noldus, the Netherlands) software for total distance travelled, and clockwise and anticlockwise rotations for each mouse. Statistical analysis was performed using one-way multivariate ANOVA implemented in SPSS24 (IBM, USA). To account for possible sex differences, age and weight were used as covariates.

### MRI setup and animal preparation

Resting-state fMRI (rsfMRI) measurements were obtained with a 7T Bruker BioSpec scanner equipped with a Pharmascan magnet and a high signal-to-noise ratio (SNR) receive-only cryogenic coil (Bruker BioSpin AG, Fällanden, Switzerland) in combination with a linearly polarized room temperature volume resonator for rf transmission.

Standardized anesthesia protocols and animal monitoring procedures were used when performing rsfMRI scans (Markicevic et al., 2020). Briefly, mice were initially anesthetized with 3% isoflurane in 1:4 O_2_ to air mixture for 3 min to allow for endotracheal intubation and tail vein cannulation. Mice were positioned on an MRI-compatible support, equipped with hot water-flowing bed to keep the temperature of the animal constant throughout the entire measurement (36.6 ± 0.5°C). The animals were fixed with ear bars and mechanically ventilated via a small animal ventilator (CWE, Ardmore, USA) at a rate of 80 breaths per minute, with 1.8 mL/min flow of isoflurane at 2%. Subsequently, a bolus containing a mixture of medetomidine (0.05 mg/kg) and pancuronium (0.25 mg/kg) was injected via the cannulated vein and isoflurane lowered to 1%. Five minutes following the bolus injection, a continuous infusion of medetomidine (0.1 mg/kg/h) and pancuronium (0.25 mg/kg/h) was started while isoflurane was further reduced to 0.5%. Animal preparation took on average 16.1 ± 2.7 minutes and all animals fully recovered within 10 min after the measurement.

### Resting-state fMRI acquisition and data preprocessing

For fMRI recordings an echo planar imaging (EPI) sequence was using with the following acquisition parameters:: repetition time TR=1 s, echo time TE=15 ms, flip angle= 60°, matrix size = 90 × 50, in-plane resolution = 0.2 × 0.2 mm^2^, number of slices = 20, slice thickness = 0.4 mm, 2280 volumes for a total scan of 38 min. Clozapine was intravenously injected 15 min after the scan start at a dose of 30 μg/kg and a total of 38 D1Cre animals (10 controls, 13 D1 excitatory and 15 D1 inhibitory mice) were scanned.

Data was preprocessed using an already established pipeline for removal of artefacts from the multivariate BOLD time-series data (Markicevic et al., 2020; Zerbi et al., 2015). Briefly, each 4D (three spatial + a temporal dimension) dataset was normalized to a study-specific EPI template (Advanced Normalization Tools, ANTs v2.1, picsl.upenn.edu/ANTS) and transferred to MELODIC (Multivariate Exploratory Linear Optimized Decomposition of Independent Components) to perform a within subject spatial-ICA with a fixed dimensionality estimation (number of components set to 60). The procedure included motion correction and in-plane smoothing with a 0.3 mm kernel. An FSL - FIX study-specific classifier, obtained from an independent dataset of 15 mice, was used to perform ‘conservative’ removal of the variance of the artefactual components (Zerbi et al., 2015). Subsequently, the dataset was despiked, band-pass filtered (0.01-0.25 Hz) based on the frequency distribution of the fMRI signal under isoflurane-medetomidine anesthesia (Grandjean et al., 2014) (Pan et al., 2013) and finally normalized into the Allen Mouse Brain Common Coordinate Framework (CCFv3) using ANTs. From each dataset, the first 900 data points (equivalent to 15 min of scanning) were used as a baseline. After the clozapine injection, we discarded the following four minutes (or 240 data points) to allow time for the DREADD to become fully activated (Markicevic et al., 2020). The subsequent 900 data points (from 19–34 min) were analyzed to estimate post-clozapine effects. The difference between baseline and post clozapine measurements is further referred to as ΔFC (for functional connectivity) or BCA (for balanced classification accuracy – see below for details).

### Defining Regions of Interest (ROI) based on structural connectivity within the striato-thalamo-cortical circuit

The Allen Mouse Brain Connectivity Atlas (AMBCA) (Oh et al., 2014) was used to map out the structural connectome of the striato-thalamo-cortical circuit containing our striatal target area CPdm. This mesoscale structural connectome of the mouse brain was derived from 469 viral microinjection experiments in the right hemisphere of C57BL/6J mice (Oh et al., 2014). The data were further annotated using the Allen Reference Atlas (Dong H. W., 2008) and summarized in a form of weighted, directed connectivity matrix with 213 brain regions estimated from a regression model (Oh et al., 2014). The model directly outputs a ‘normalized connection strength’, estimated by scaling the injection volume in a source region to explain the segmented projection volume in a target region, and a *p*-value for each edge in the connectome. This was used to construct a 213 × 213 ipsilateral connectivity matrix. For the purposes of this study, we used the edges with the greatest statistical evidence (*p*-value < 0.05) from the ipsilateral connectivity matrix to estimate regions involved in the striato-thalamo-cortical circuit.

To estimate which regions of interest (ROIs) form the striato-thalamo-cortical loop, including CPdm, we took a stepwise approach. First, starting from the caudate putamen (CP) as our “seed area” we identified globus pallidus external (GPe)/ internal (GPi) and substantia nigra pars compacta (SNc)/pars reticulata (SNr) as regions that CP directly projects to. Second, GPe/GPi and SNc/SNr were used as seed areas to identify subthalamic nuclei (STN) and several thalamic ROIs as being directly connected (RT, PP, LH, VM, PF, VAL, IMD, SPA, SPFp, VPMpc and POL, see the caption of Fig. 1 for definitions of the abbreviations). Third, using the identified thalamic regions as a seed, we found direct projections to other thalamic subregions (MD, CM, AM, VPM, VPL, PO, CL, SPFm, LP, RH, RE, LD; cf. Fig. 1). Fourth, using all thalamic ROIs, which were identified in the second and third steps, we identified connected cortical ROIs that also have significant projections to CP, thereby closing the loop. These cortical regions were ACA, AI, MOp/s, SSp, GU, VISC, PL, RSP, ORB, PERI, VIS, PTLp (cf. Fig. 1).

Some of the extracted thalamic and cortical ROIs consisted of only a few voxels and were consequently merged to improve the signal-to-noise ratio for the BOLD time-series analyses. The criteria for merging ROIs were that they: (i) were anatomically located next to each other; and (ii) had similar anatomical connectivity patterns derived from Allen Brain atlas. The following groups of thalamic ROIs were merged: 1) RE, LH and RH, 2) SPF, SPA and PP, 3) PO and POL, 4) VAL, VPM and VPMpc, 5) VM and CM; and the following cortical ROIs were merged: 6) VISC and GU and 7) ECT and PERI. This resulted in 14 thalamic (TH) ROIs and 12 cortical (CTX) ROIs.

The mesoscale structural connectome (Oh et al., 2014) contains the caudate-putamen (CP) as a single area. Since we neuromodulated the dorsomedial subarea of CP, we further refined the striatal parcellation using more specific anatomical information. Recent research has shown that the CP consists of functionally segregated parts (Hintiryan et al., 2016) that can be distinguished along a rostral-caudal gradient, a dorsal-ventral gradient, and a medial-lateral gradient. Using this information from the Mouse Cortico-Striatal Projectome (Hintiryan et al., 2016), we further parcellated caudate-putamen into 29 distinct regions.

Finally, we used information from the literature (Alloway et al., 2017; Collins et al., 2018; Diaz-Hernandez et al., 2018; Evangelio et al., 2018; Guo et al., 2015; Hunnicutt et al., 2016; Lee et al., 2020; Mandelbaum et al., 2019; Parker et al., 2016; Perry & Mitchell, 2019; Wall et al., 2013) to identify which of the thalamic nuclei from the structural connectome identified above (Oh et al., 2014) were specifically anatomically connected to CPdm (our targeted subarea of the caudate putamen). Based on this information, we refined our selection of thalamic ROIs to VM|CM (|-indicates merging of these ROIs) , LD, MD, PF, LP, and CL, which are all reciprocally connected to CPdm, i.e., these thalamic nuclei receive projections from CPdm via the GP/SN and project back to the CPdm (Suppl. Table 2). In summary, the striato-thalamo-cortical circuit consisted of 14 cortical areas (CTX), 21 thalamic areas (TH), 29 striatal subareas, globus pallidus internal and external (GPi/e), substantia nigra pars reticulata and pars compacta (SNr/c), and subthalamic nucleus (STN).

### Resting-state fMRI data analysis

#### Classifying univariate BOLD time series

To understand how cellular-level manipulations shape the BOLD dynamics of structurally well-defined striato-thalamo-cortical circuitry at the macroscale, time series of each ROI in this circuit were obtained. Each univariate BOLD time series was represented as a large feature vector, where each feature corresponds to an interpretable summary statistic computed from the time series. Feature extraction was performed using the *hctsa* toolbox v1.04 (Fulcher & Jones, 2017; Fulcher et al., 2013), which computes 7702 time-series features per time series. For each ROI, features were computed for each of two time periods (baseline vs post clozapine injection) in each of 38 individual subjects (*n*=13 D1 excitation, *n*=15 D1 inhibition, and *n*=10 controls). We filtered out features which were poorly behaved due to having constant values or non-real-valued feature outputs, across time series within all three groups and for each ROI separately.

Feature-based representations of BOLD dynamics in each brain area were used as the basis for classifying different experimental conditions. Because our analysis focused on *changes* of time series features due to DREADD activation, we subtracted time series features computed at baseline from those obtained during the post-clozapine period. To classify a given brain area’s BOLD dynamics, the subtracted features were normalized using an outlier-robust sigmoidal transformation (Fulcher et al., 2013) and used as the basis for classification using a linear support vector machine (SVM). To account for class imbalance (n_control_ = 10; n_excitation/inhibition_ = 13/15), we used inverse probability class reweighting to train the SVM, and balanced accuracy (the arithmetic mean of sensitivity and specificity) to evaluate performance. Classification performance was assessed as the mean 10-fold stratified cross-validated balanced accuracy (BCA – balanced classification accuracy), averaged over 50 repetitions (to reduce variance caused by the random partition of data into 10 folds).

In small samples, there is a greater probability that false-positive classification results can be obtained by chance. To account for this effect, the statistical significance of each classification result was estimated using a permutation test. We permuted the assignment of group labels to time series to construct a null distribution of the BCA metric for each region (mean across 50 repeats of 10-fold mean cross-validated BCA) for 5000 random group-label assignments to the data. This allowed us to estimate a *p*-value for each region, which was corrected across all regions by controlling the false discovery rate (FDR) using the method of Benjamini and Hochberg (Benjamini & Hochberg, 1995). For individual regions with significant classification performance, we next aimed to understand which, if any, individual features were discriminative of D1 excitation from controls. To this end, we scored the discriminability of each feature using a Mann-Whitney rank-sum test and computed corrected *p*-values across all the 6480 features using FDR correction for each ROI separately.

#### Resting-state fMRI functional connectivity analysis

Functional connectivity between CPdm and all other ROIs was measured using regularized Pearson’s correlation coefficients calculated in FSLnets. A one-sample Wilcoxon test was performed to test whether baseline functional connectivity differs from 0 (Fig. 7A). For each ROI, normalized (post-clozapine – baseline) connectivity values were entered into a General Linear Model (GLM) implemented in FSL and pairwise comparisons among the 3 groups (D1 excitation, D1 inhibition, and D1 control) performed. Age and weight were used as covariates. Permutation testing with 5000 permutations was performed to estimate whether DREADD activation caused functional connectivity changes between groups, followed by the correction for multiple comparisons across all the regions using FDR with a significance threshold, *p* < 0.05. To visualize DREADD-induced changes in FC, baseline connectivity was subtracted from post clozapine connectivity values, thus ΔFC > 0 indicates a DREADD-induced increase in FC and ΔFC < 0 indicates a decrease.

#### Histological evaluation of transfection

DREADDs viral expression (for DIO-hM3Dq-mCherry, DIO-hM4Di-mCherry, and DIO-mCherry) was confirmed by mCherry staining using standard immunohistochemistry protocols, while qualitative transfection of D1 MSNs was confirmed using antibodies against D1 marker prodynorphin. Briefly, after the last MRI session, mice were deeply anesthetized using a mixture of Ketamine (100 mg/kg; Graeub, Switzerland), Xylazine (10 mg/kg; Rompun, Bayer) and Acepromazine (2 mg/kg; Fatro S.p.A, Italy) and transcardially perfused with 4% *Paraformaldehyde* (PFA, pH=7.4). The brains were postfixed in 4% PFA for 1.5 hours at 4°C and then placed overnight in 30% sucrose solution. Brains were frozen in a tissue mounting fluid (Tissue-Tek O.C.T Compund, Sakura Finetek Europe B.V., Netherlands) and sectioned coronally in 40 μm thick slices using a cryostat (MICROM HM 560, histocom AG-Switzerland). Free-floating slices were first permeabilized in 0.2% Triton X-100 for 30 min and then incubated overnight in 0.2% Triton X-100, 2% normal goat serum, guinea pig anti-prodynorphin (1:500, Ab10280, Abcam) and rabbit anti-mCherry (1:1000, Ab167453, Abcam) or rabbit anti-cfos (1:5000, AB2231974, Synaptic systems) at 4°C under continuous agitation (100 rpm). The next day, sections were incubated for 1h in 0.2% Triton X-100, 2% normal goat serum, goat anti-rabbit Alexa Flour 546 (1:300, A11035, Life Technologies), goat anti-guinea pig Alexa Flour 647 (1:200, cat #A-21450, ThermoFisher Scientific) and DAPI (1:300, Sigma-Aldrich) at room temperature under continuous agitation. Afterwards, slices were mounted on the superfrost slides where they were left to airdry and later coverslipped with Dako Flourescence mounting medium (Agilent Technologies). A confocal laser-scanning microscope (CLSM 880, Carl Zeiss AG, Germany) and Zeiss Brightfield microscope (Carl Zeiss, AG Germany) were used to detect the viral expression. The microscopy protocol included a tile scan with a 10x or 20x objective, pixel size of 1.2 μm and image size of 1024×1024 pixels. Images were preprocessed and analyzed using ImageJ-Fiji.

## Acknowledgements

We thank the team of the EPIC animal facility for providing animal care. We thank Jean-Charles Paterna from the Viral Vector Facility (VVF) of the Neuroscience Center Zurich, a joint competence center of ETH Zurich and University of Zurich for producing viral vectors and viral vector plasmids. This work was supported by ETH Research (grant ETH-38 16-2 to N.W.), SNSF AMBIZIONE (PZ00P3_173984/1 to V.Z.).

## Author Contributions

M.M. designed the study, performed the experiments and analyses, and wrote the paper. O.S. analyzed behavioral videos. J.B. provided supervision and feedback on behavioral study design and the paper. M.R. provided supervision of fMRI experiments and feedback on the paper. V.Z., B.D.F. and N.W. provided supervision and feedback on the study design and analyses and wrote the paper.

## Supplementary Figures

**Suppl. Figure 1:**
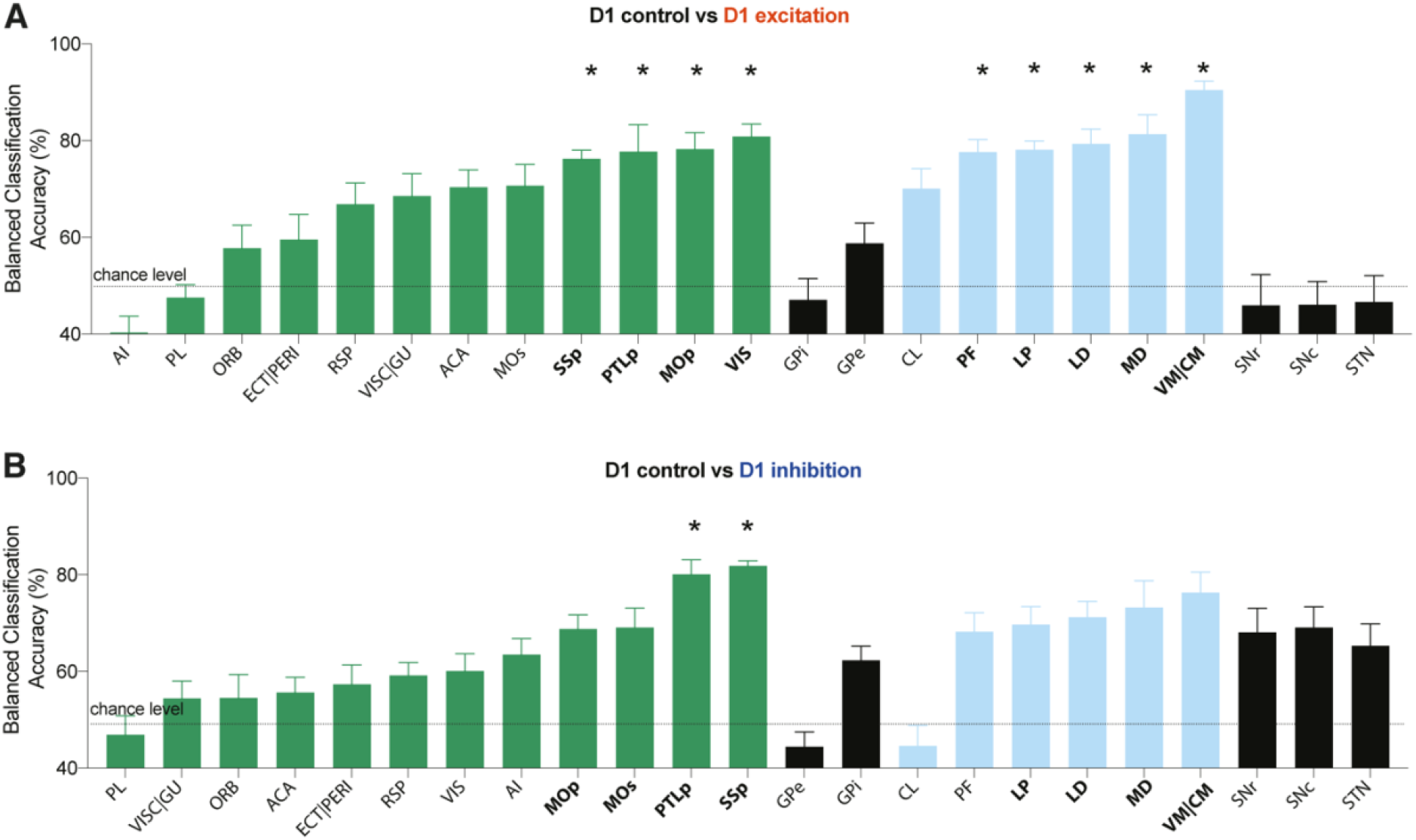
Characteristic changes observed in BOLD dynamics of regions part of the striato-thalamo-cortical circuit. Classification results in each brain region are shown for **A)** D1 control versus D1 excitation; **B)** D1 control versus D1 inhibition. Each region is color coded depending on whether it is a part of CTX, TH or other regions downstream of basal ganglia. A significant differences in BOLD dynamics between the two groups (permutation test, uncorrected, p<0.05) is depicted with a bolded ROI abbreviation, where *indicates that statistical significance was reached after FDR correction for multiple comparison.

## Supplementary Tables

**Suppl. Table 1:**
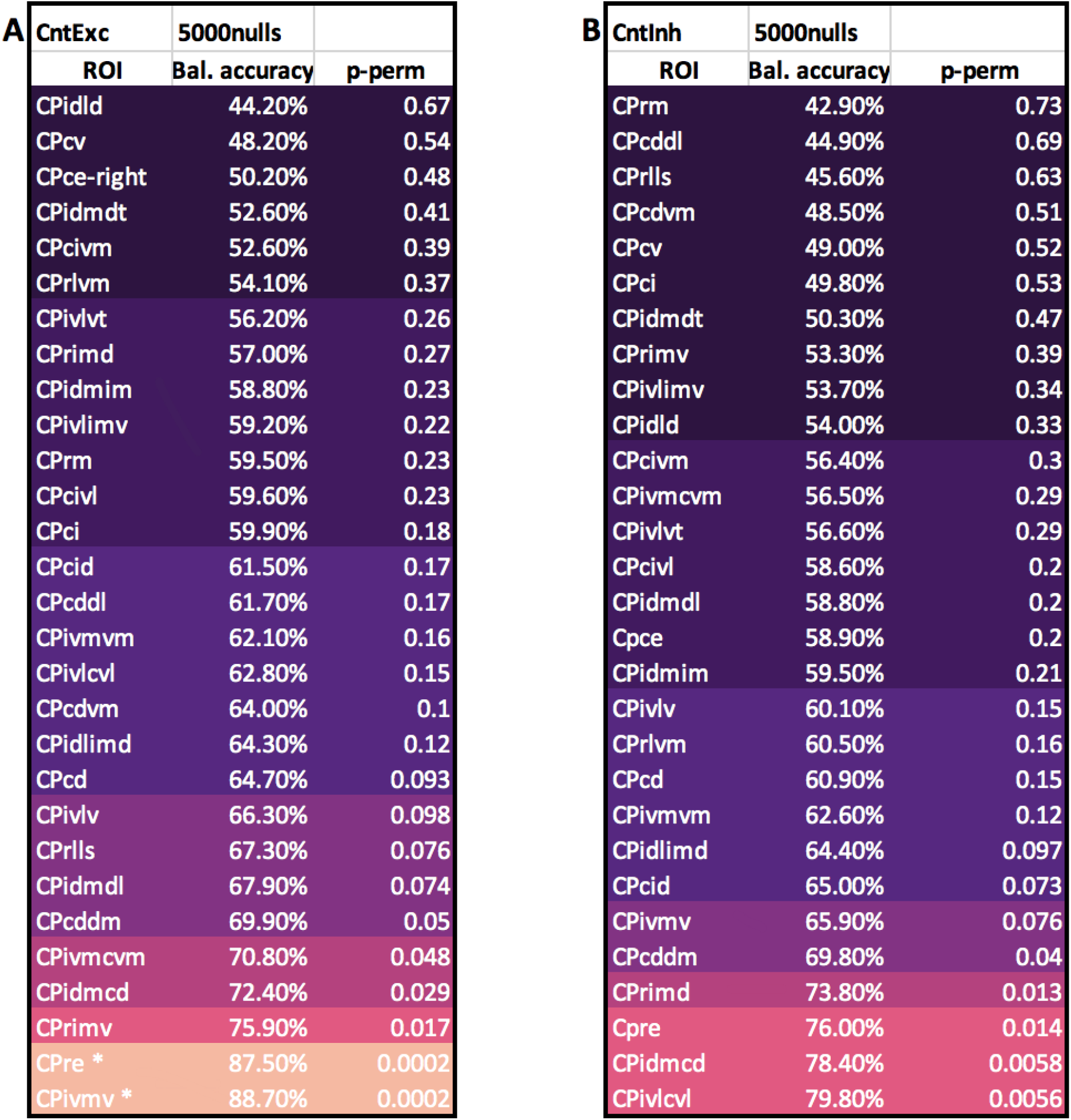
Balanced accuracies for 29 subregions of CP obtained from Mouse-Striatal Projectome. **A)** Balanced classification accuracy results for a pair of groups i.e., D1 control vs D1 excitation together with permutation test p-values calculated using 5000 nulls. All p-values are uncorrected for multiple comparisons. * indicates ROIs survived the FDR correction for multiple comparisons (p<0.05). **B)** Identical to A but for D1 control vs D1 inhibition. The exact location of ROIs with balanced accuracies are illustrated in Fig. 3A-B, respectively.

**Suppl. Table 2:** List of thalamic ROIs and their anatomical projections from/to dorsomedial CP with references to the literature where the information was obtained.

| Thalamic nuclei | Projections **FROM** CPidm (via Gpi/SNr) | Projections **TO** CPidm | Reference(s) |
| --- | --- | --- | --- |
| CL | yes | yes | Parker et al., 2016; Wall et al., 2013; Guo et al., 2015; Kamishina et al., 2008 |
| CM | yes | yes | Wall et al., 2013; Parker et al., 2016; |
| PF | yes | yes | Mandelbaum et al., 2019; Lee et al., 2020; Wall et al., 2013 |
| MD | yes | yes | Hunnicutt et al., 2016; Collins et al., 2018; Wall et al., 2013 |
| LP | yes | yes | Guo et al., 2015; Evangelio et al., 2018; Kamishina et al., 2008; Alloway et al., 2017; |
| LD | yes | yes | Hunnicutt et al., 2016; Perry & Mitchell, 2019; |
| VM | yes | yes | Lee et al., 2020; Wall et al., 2013; Collins et al., 2018 |
| RE | no | no | Vertes et al., 2016; |
| LH | no | no | Namboodiri et al., 2016 |
| RH | no | yes (to dorsal CP but scarce) | Parker et al., 2016; Vertes et al., 2016 |
| PO | no | no | El-Boustani et al., 2020; Kamishina et al., 2008; Alloway et al., 2017; |
| POL | no | no | El-Boustani et al., 2020; Alloway et al., 2017 |
| IMD | no | no | Van der Werf et al., 2002; Smith et al., 2004 |
| RT | no | no | Antal et al., 2014; Hunnicutt et al., 2016; |
| AM | no | yes (to dorsal CP but scarce) | Perry & Mitchell, 2019; Wall et al., 2013; Bubb et al., 2017 |
| VPL | no | no/scarce | Antal et al., 2014; Diaz-Hernandes et al., 2018; Erro et al., 2002 |
| VPM/VPMpc | no/scarce | no | Hunnicutt et al., 2016; Diaz-Hernandes et al., 2018 |
| VAL | no | no | Alloway et al., 2017; Smith et al., 2004; |
| SPF | no | no | Palkovits et al., 2006; Oh et al., 2014 |
| SPA | no | no | Palkovits et al., 2006; Oh et al., 2014 |
| PP | no | no | Linke et al., 2000; Oh et al., 2014 |

